# Ultra-low-field paediatric MRI in low- and middle-income countries: super-resolution using a multi-orientation U-Net

**DOI:** 10.1101/2024.02.16.580639

**Authors:** Levente Baljer, Yiqi Zhang, Niall J Bourke, Kirsten A Donald, Layla E Bradford, Jessica E Ringshaw, Simone R Williams, Sean CL Deoni, Steven CR Williams, Khula SA Study Team, František Váša, Rosalyn J Moran

## Abstract

Owing to the high cost of modern MRI systems, their use in clinical care and neurodevelopmental research is limited to hospitals and universities in high income countries. Ultra-low-field systems with significantly lower scanning costs present a promising avenue towards global MRI accessibility, however their reduced SNR compared to 1.5 or 3T systems limits their applicability for research and clinical use. In this paper, we describe a deep learning-based super-resolution approach to generate high-resolution isotropic T_2_-weighted scans from low-resolution paediatric input scans. We train a ‘multi-orientation U-Net’, which uses multiple low-resolution anisotropic images acquired in orthogonal orientations to construct a super-resolved output. Our approach exhibits improved quality of outputs compared to current state-of-the-art methods for super-resolution of ultra-low-field scans in paediatric populations. Crucially for paediatric development, our approach improves reconstruction of deep brain structures with the greatest improvement in volume estimates of the caudate, where our model improves upon the state-of-the-art in: linear correlation (r = 0.94 vs 0.84 using existing methods), exact agreement (Lin’s concordance correlation = 0.94 vs 0.80) and mean error (0.05 cm^3^ vs 0.36 cm^3^). Our research serves as proof-of-principle of the viability of training deep-learning based super-resolution models for use in neurodevelopmental research and presents the first model trained exclusively on paired ultra-low-field and high-field data from infants.

## 1 Introduction

Neuroimaging studies on infant and child populations have become increasingly vital in establishing the relationship between neurodevelopment and resultant cognitive functioning. Magnetic resonance imaging (MRI) has proven to be particularly essential in this endeavour, owing to its ability to provide insight into structural, functional and metabolic brain development, in addition to revealing pathologies pertaining to neurobiological disorders (Nolen-Hoeksema et al., 2014). Accessing these capabilities, however, is only possible by overcoming significant financial barriers: on top of the cost of a scanner itself (roughly $1,000,000 per Tesla; Arnold et al., 2023), the high field strength of existing systems requires facilities with electromagnetic shielding and specialised staffing. Furthermore, since most modern MRI systems use superconducting magnets, they require the use of cryogens, which themselves come with storage, transportation and maintenance costs (Sarracanie et al., 2015). Altogether, these expenses establish strict financial boundaries on neuroimaging studies and clinical work. Even in hospitals and universities within high-income countries (HICs), scanning costs ($500-$1000/hr per research scan) place a limit on the number of participants scanned and the duration of longitudinal research.

More concerningly from a global health perspective, infrastructural costs severely limit MRI accessibility in low- and middle-income countries (LMICs). Evaluating MRI accessibility based on ratio of MRI units per million population, HICs such as Japan and the US boast 51.67 and 38.96 units/million, respectively (Ogbole et al., 2018). Comparing these values with those of Nigeria and Ghana – which own 0.30 and 0.48 units/million, respectively – reflects a hundred-fold difference between HICs and LMICs (Jalloul et al., 2023). Consequently, our current understanding of neurodevelopment in such regions, where adversities pertaining to nutrition, sanitation and higher rates of infectious risk play a crucial role, is primarily based on psychometric measures or low-cost, functional imaging methods such as EEG or fNIRS (Perdue et al., 2019, Jensen et al., 2021).

Ultra-low-field (ULF) imaging systems with magnetic field strengths ranging from 50 to 100mT (e.g., the 64mT Hyperfine Swoop) present a potential solution to issues of access inequality (Sarracanie et al., 2015, Abate et al., 2024). Such systems rely on the use of large, permanent magnets instead of superconducting electromagnets and as such have significantly reduced component prices, reduced room requirements, and reduced costs for power, cooling and maintenance compared to high-field (HF) imaging systems (Campbell-Washburn et al., 2019). Unfortunately, these benefits are paired with a significantly diminished signal-to-noise ratio (SNR) (Klein, 2020), rendering image outputs sub-optimal for visual reading by radiologists or for a large portion of automated processing methods in currently available neuroimaging toolkits. Such processing methods usually require high-resolution MRI scans (1mm isotropic), whereas product sequences on the Hyperfine system yield a default spatial resolution of 1.5×1.5×5 mm in 3-6 minutes per image. Higher resolution images may be acquired with increased acquisition times (∼12–15 min for a single 2×2×2mm T_2_-weighted image), although this may decrease patient compliance and satisfaction and increase the risk of head motion, particularly in sensitive populations such as infants or elderly participants (Madan, 2018; Padormo et al., 2023).

One technique that has found success in enhancing the SNR of ULF outputs is super-resolution (SR) reconstruction of a single image from multiple lower-resolution images, acquired in three orthogonal orientations (i.e., axial, sagittal, and coronal) (Deoni et al., 2022). This is carried out through repeated multi-resolution registration (MRR) of the scans using the Advanced Normalization Tools (ANTs) multivariate template construction command. Here, low-resolution images are aligned using linear and diffeomorphic registration with symmetric normalization, outputting a final image with effective dimensions of 1.5×1.5×1.5mm (Figure 1). Although this approach still entails a longer scanning session (up to ∼18 min to acquire three scans), multiple shorter acquisitions may reduce the risk of head motion within each scan compared to a single higher-resolution acquisition. Furthermore, a single instance of head motion would only corrupt one of three scans instead of the whole higher-resolution scan.

**Figure 1).**
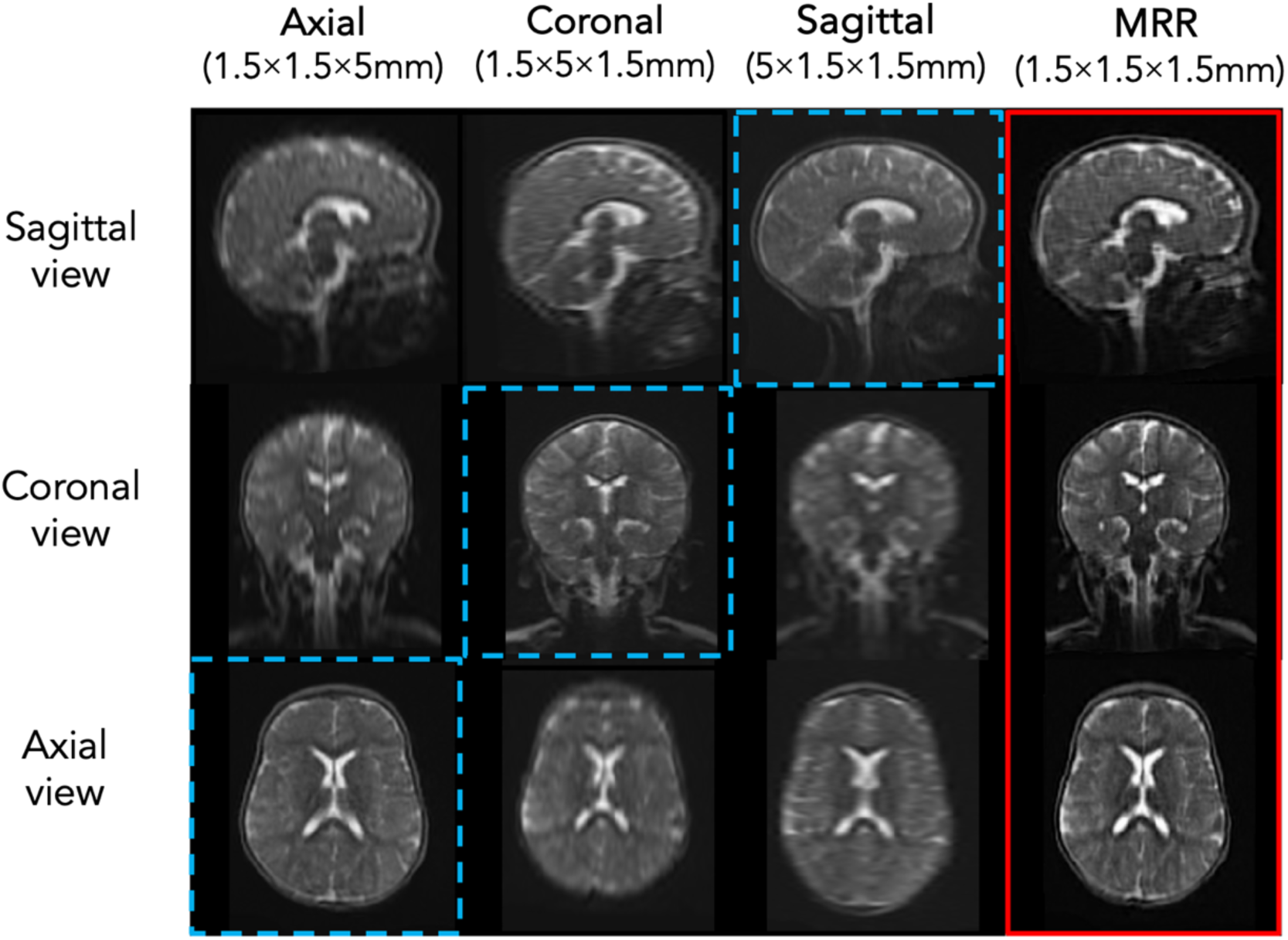
Columns 1-3: non-isotropic Hyperfine scans of a 6-month-old subject (high-resolution plane highlighted with dashed blue lines); Column 4: isotropic output of MRR (highlighted in red)

An alternative approach involves the use of modern image translation methods based on convolutional neural networks (CNNs), which learn a transformation between a source image and a target image. A decade of research has gone into the use of CNNs for super-resolution, with architectures ranging in complexity from simple three-layer models (Dong et al., 2014) to twenty-layer models (Kim et al., 2016). In models tailored for medical image super-resolution, a similar variety in complexity and functionality is present (Oktay et al., 2016), (Pham et al., 2019), however the U-Net (Ronneberger et al., 2015) has consistently presented itself as one of the most reliable architectures for use in a wide array of MRI image translation tasks (Kelly et al., 2022). The architecture consists of a contracting path to capture context and a symmetric expanding path that enables precise localization of biological structures. Through the contracting path, spatial information is normally lost, however skip connections between the two paths ensure feature reusability.

In their standard form, super-resolution U-Nets train to minimise a voxel-wise loss between input and target images (such as L1 or L2). This ensures appropriate outputs by constraining voxel values of generated high-resolution images to be close to those of the ground truth. Unfortunately, such losses do not take perceptual quality or textures into account, resulting in outputs that are often perceptually unsatisfying and lack high frequency details (Wang, 2019; Zhang, 2018). As such, more recent super-resolution models tend to supplement traditional voxel-wise loss functions via the addition of other losses, for instance a Dice loss (Iglesias et al., 2021, 2023) or an adversarial loss (Zhou, 2022).

In our paper, we employ an alternative loss function that relies on the internal activations of deep CNNs trained on high-level image classification tasks. The computer vision community has noted the effectiveness of such deep CNNs and related features in their correspondence to human perceptual judgments (Zhang, 2018). As such, by adding a loss term targeted on extracted image features, we can encourage feature similarity between predicted and real images and, in turn, enhance perceptual similarity. Unlike a Dice loss, this operates directly on the level of image features, and unlike an adversarial loss, it allows us to improve perceptual quality without the computational burden of training a classifier in addition to our generator. The latter factor is crucial for our training needs, for it frees us to feed three separate 3D volumes into the generator at each training step, as inspired by the MRR approach of combining three ULF scans into a single high-resolution output. Here we present the Multi-Orientation (MO) U-Net, which produces a synthetic 1mm isotropic scan from three non-isotropic ULF input scans. To the best of our knowledge, deep learning-based SR of ultra-low-field brain MRI has only been investigated in adult populations, where in many cases models are being trained on partially or even entirely synthetic datasets. As such, we propose the first U-Net trained on real, paired ULF-HF data from a paediatric population, and demonstrate its ability to surpass alternative SR techniques.

## 2 Methods

### 2.1 Imaging data

MRI data used in this paper stems from a study investigating the neurodevelopment of executive function over the first 3 years of life in participants based in South Africa and Malawi (Zieff et al., 2024, Abate et al., 2024). Images used here included subjects from South Africa exclusively, at ages of either 3- or 6-months, with no known neurological abnormalities. A total of 82 subjects attended two scanning sessions (ULF and HF); all subjects had HF T_2_w scans acquired using a Siemens 3T scanner (Erlangen, DE) and had ULF T_2_w scans acquired using a Hyperfine Swoop 64mT system (Guildford, CT), with high in-plane resolution along three orthogonal planes (axial, sagittal, coronal). Of these subjects, 63 successfully completed all four scanning protocols (one HF and three ULF scans). Pre-processing involved rigid registration between all HF scans and a custom age-specific HF template, and each subject’s ULF with the corresponding pre-registered HF scan (Ashburner, 2007). Following this, we skull-stripped HF and ULF scans separately using HD-BET, a deep-learning tool for MRI brain extraction (Isensee et al., 2019). Seven subjects were excluded due to major artifacts being present in at least one of their ULF scans (see supplementary Figure S1 for details), thus our final sample size consisted of N*=*56 subjects (26 Male, 30 Female), of which 16 were scanned at 3-months and 40 at 6-months. Each subject had three ULF (64mT; 1.5×1.5×5 mm) and one HF (3T; 1×1×1×mm) T_2_-weighted scans.

To maximise the data available from a single subject, we fed all three ULF scans into our network as input (Hyperfine scans acquired in three orthogonal orientations), thus each matched ‘pair’ of ULF-HF scans included three orthogonal ULF scans and one HF scan. We split these pairs across 4 folds with a training/validation/test split of 42/7/7. As such, 7 subjects were used in each fold to monitor validation loss, and inference was carried out on a total of 28 subjects (7 subjects x 4 folds). Age and gender stratification was applied for each split, across each of the 4 folds (see supplementary Table S1 for demographic details on splits).

### 2.2 CNN architecture and training protocol

Our MO U-Net builds on the architecture of the 3D U-Net (Çiçek et al., 2016). It has three input channels jointly flowing into three encoding levels, each consisting of a convolution with a 3×3×3 kernel (allowing the model to capture fine-grained details while reducing parameter requirements), group normalisation (speeding up convergence by preventing internal covariate shift), a rectified-linear unit (ReLU) activation and maxpooling. The first layer has 64 features, with the number of features doubling after each maxpooling and halving after each upsampling. The final layer uses a linear activation to produce the final image output. All input and output images were resized (and padded where necessary) to have a uniform size of 160, 160, 160. The model is implemented using PyTorch.

Our MO U-Net was trained across all 4 folds until convergence (1500-2000 epochs) using the Adam optimizer (Kingma, 2014) with a learning rate of 10^−4^. The average training time was 41.2 hours using an Nvidia RTX 3090 GPU. ULF images were normalised and scaled to 1-mm isotropic resolution, ensuring that input and output resolutions match. Augmentation of images was carried out on the fly, with random affine transform or random elastic deformation applied to both ULF and HF data with pre-set probability (p = 0.5). Batch size was set to 1, as feeding in more than one triplet of ULF images to our model at each training step exceeded our GPU capacity. After training, weights of the model were frozen and inference was carried out (∼1 second on an Nvidia RTX 3090 GPU and ∼30 seconds on a modern CPU).

### 2.3 Loss function

To capture anatomical similarity between predicted and ground truth images, the training of our network minimises the L2-norm with respect to the neural network parameters θ:

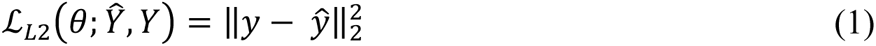

where ***ŷ*** denotes the model output and ***y*** denotes the target image. Although necessary for pair-wise image-translation, previous work has noted that training exclusively using a voxel-wise loss can diminish the quality of outputs and result in excessively smooth SR images (Zhang et al., 2022). We support this finding with our own analyses (see supplementary Table S2 and Figure S2). As such, we add a perceptual loss to enhance similarity in features between input and target images by employing the Learned Perceptual Image Patch Similarity (LPIPS) (Zhang et al., 2018). This metric quantifies the similarity between features from two separate images, as extracted by a pre-trained classifier. For this, we use AlexNet (Krizhevsky et al., 2012). Although other classifiers can be used for feature extraction, including the much larger VGG (Simonyan et al., 2014), we employ AlexNet it allows for more efficient training and has been shown to provide deep embeddings which agree equally well with humans (Zhang et al., 2018). The formula for this distance metrics is shown below:

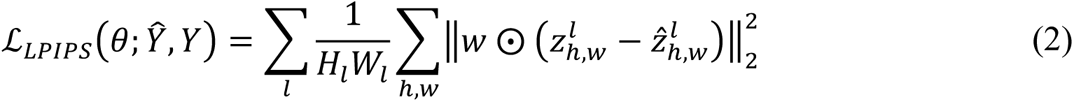

where ***ẑ*** denotes the features of prediction ***ŷ*** and ***z*** denotes the features of ground-truth ***y*** extracted from the first 5 layers of AlexNet, with ***ẑ^l^****, **z^l^*** ∈ ℝ^HxWxC^ for each layer ***l***. Features ***ẑ*** and ***z*** are first normalised across the channel dimension then scaled across the same dimension by ***w^l^*** ∈ ℝ^C^ (a hyperparameter set by the original authors of LPIPS), after which an L2 norm is computed between the two values. The difference is averaged spatially then summed across layers to produce the final output. To train our MO U-Net, we combine equations 1 & 2, yielding the following loss:

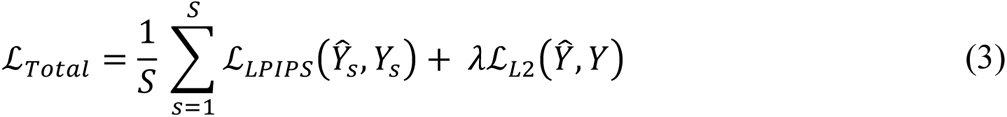

where ***λ*** is a hyperparameter set at ***λ***=100, inspired by previous image-to-image translation work (Isola et al., 2016). Importantly, AlexNet was trained to classify 2D images, therefore random, paired slices ***Ŷ_s_*** and ***Y_s_*** are taken from the set of predictions and ground-truth images *Ŷ* and *Y* to compute an average LPIPS score across all slices ***S***. To allow for multiple slices to be assessed across all volumetric dimensions while minimising the number of computations needed to be performed at each training step, we set ***S*** to 6. The full training scheme is depicted in Figure 2.

**Figure 2).**
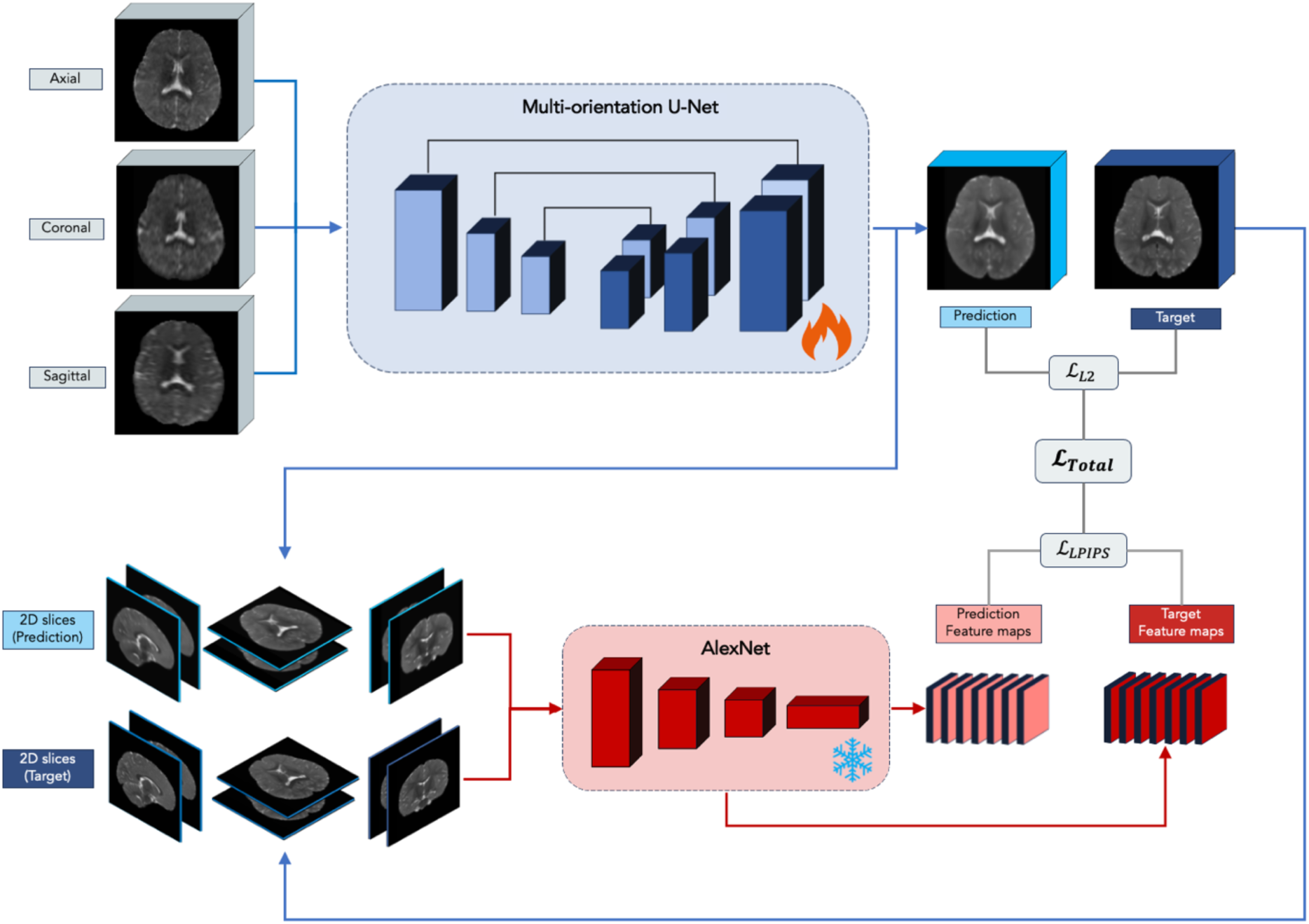
Model training: Multi-orientation U-Net (in blue) with three input channels and one output channel, allowing a transformation from three ULF scans to one SR image. Outputs of this model, along with ground-truth high-field images, are fed into a frozen AlexNet for feature extraction. The final loss is calculated by comparing model outputs with corresponding high-field scans, both at the level of full images and their feature space

### 2.4 Related work and model evaluation

Although the literature surrounding SR of medical images is vast, research centered specifically on SR of ULF scans is comparatively scarce, largely owing to the relatively recent emergence of such imaging systems. As discussed in the introduction, the use of multi-resolution registration by Deoni et al. (2022) stands as the state-of-the-art in the space of non-deep learning-based SR of ULF scans. SynthSR emerged as the first publicly available CNN-based SR toolkit for use on images of any contrast and resolution (Iglesias et al., 2021), with a subsequent dedicated ultra-low-field model that transforms non-isotropic T_1_w or T_2_w images to 1mm isotropic T_1_w scans (Iglesias et al., 2023). LoHiResGAN soon followed, introducing a model based on the Pix2Pix architecture to generate synthetic 1mm isotropic scans from non-isotropic inputs acquired at 64mT (similar to our model) (Islam, 2023). Additional work was also carried out on alternative resolutions and field strengths using a model based on multiscale feature extraction and spatial attention (Lau, 2023). This was used to generate synthetic 1.5mm isotropic scans from 3mm isotropic inputs acquired at 55mT (Man, 2023) and 1mm isotropic scans from 3mm isotropic inputs acquired at 50mT (Zhao, 2024). For comparative analyses, we restrict ourselves to techniques designed for an identical image translation task as ours (MRR) or of any field-strength and resolution (SynthSR version 2.0).

As such, in addition to assessing the extent to which our MO U-Net enhances ULF inputs, we compared its performance against MRR and SynthSR using the metrics described below:

#### 2.4.1 Segmentation-based metrics

The quality of model outputs relative to ground-truth HF images was evaluated via Dice overlap of segmented brain regions within subjects and tissue volume correlations across subjects. Both segmentation outputs and volume estimations were obtained using SynthSeg (Billot, 2023), a segmentation toolkit that is agnostic to contrast and resolution. This feature of SynthSeg allowed us to include SynthSR in our comparative analyses, which exclusively outputs T_1_w scans regardless of the contrast of the input. In addition to segmenting cortical grey matter (GMC), subcortical grey matter (GMS), white matter (WM) and cerebrospinal fluid (CSF), we segmented the following deep brain structures: accumbens, amygdala, pallidum, hippocampus, caudate, putamen, thalamus, and ventral diencephalon. Considering that one of the primary applications of successful SR techniques would be use in ULF-based neurodevelopmental research in LMIC settings, selecting regions that play an important role in neurodevelopmental outcomes is particularly beneficial to underscore the value of our technique. These deep grey matter nuclei are known to be affected in conditions such as hypoxic-ischemic encephalopathy (HIE) (Hassett et al., 2022) or perinatal stroke (Ilves et al., 2022), such that damage is associated with worse neurodevelopmental outcomes. All segmentations were visually inspected prior to running analyses, with two test subjects being removed due to failed segmentations (see supplementary Figure S3 for more details on the segmentations and supplementary Table S3 for demographics after exclusion). As such, a final sample of N=26 subjects was used for segmentation-based analyses.

#### 2.4.2 Intensity differentiation and image quality assessment

We additionally assessed model performance by directly comparing the output images. We first investigated whether the separation between grey matter and white matter was enhanced via our MO U-Net, based on the difference in intensities between these two tissue types. This was done by applying grey matter and white matter segmentations from high-field scans as masks to other images (ULF scans, MRR outputs, MO U-Net outputs and HF scans), followed by calculating the difference in median voxel intensities between these two regions. Since this metric was used to assess whether GM and WM separation in super-resolved images more closely matched that of high-field T_2_w scans, the metric was not calculated for SynthSR.

Image quality was additionally evaluated using normalized mean squared error (NMSE), peak signal-to-noise ratio (PSNR), and structural similarity index (SSIM) between predicted images and corresponding high-field scans. Once again, the T_1_w SynthSR outputs could not be directly compared to the T_2_w outputs of MRR or MO U-Net. Nevertheless, we were able to calculate these metrics for SynthSR outputs on a subset of the 28 test subjects who also had a T_1_w 3T scan acquired as part of their scanning protocol (N=25). See supplementary Table S3 for details on the demographic distribution within this subset.

#### 2.4.3 Performance with reduced input

Using our trained MO U-Net, we additionally assessed performance with varying numbers of input scans. More specifically, we assessed model outputs when the MO U-Net only received an axial scan or axial and sagittal scans (as opposed to all three orientations, with which it was originally trained). Since our trained model requires three input images, inference with missing scans was achieved by cloning the axial scan either once or twice, depending on whether one or two distinct inputs were provided. As such, we ran inference with each subject a total of three times, with the following combination of scans: 1) axial, axial, axial; 2) axial, axial, sagittal; 3) axial, sagittal, coronal. The choice of prioritising scans in the order of axial > sagittal > coronal was done based on the scanning protocol used, which involved scanning participants in this order.

### 2.5 Statistical methods

We compared the within-subject Dice overlap of segmentations between pairs of models using the Wilcoxon signed-rank tests to compare the medians (Wilcoxon, 1945). Tissue volume correlations across participants were assessed using both Pearson’s correlation coefficient (Pearson, 1895), determining linear correlation, and Lin’s concordance correlation coefficient (Lin, 1989), determining exact agreement, or alignment with the x = y identity line. Accuracy of volume estimations was additionally assessed by reporting the mean difference in volumes obtained from high-field segmentations and SR outputs. The choice of analyses is based on previous work investigating correspondence between ULF and HF scanners (Váša et al., 2024).

## 3 Results

### 3.1 Segmentations

Segmentations obtained from MO U-Net predictions significantly improved compared to those obtained from ULF scans (Figure 3). This is highlighted by an increased Dice overlap of segmented MO U-Net predictions and HF scans compared to Dice overlap of segmented ULF and HF scans across all brain four global tissue types (Figure 4). Comparing Dice overlap of model predictions and HF scans to that of MRR outputs and HF scans, median Dice scores increased on average by 0.023. The greatest difference was seen in subcortical grey matter (Δ Dice = 0.034) and the lowest in white matter (Δ Dice = 0.015). In comparison to SynthSR outputs, median Dice scores of MO U-Net outputs rose on average by 0.067, with the greatest difference in CSF and lowest in white matter (Δ Dice = 0.138 and 0.007, respectively). Across age groups, we note that Dice overlap of all images assessed was higher in 6-month-old subjects than 3-month-old subjects (see supplementary Table S4). Significance testing using the Wilcoxon signed-rank test revealed significant improvement in MO U-Net Dice scores for all global tissue types relative to MRR, and all tissue types except subcortical grey matter (GM) when compared to SynthSR, given a family-wise error rate (FWER) of 0.00625 (Table 1). Furthermore, volume-based analysis of global tissue types revealed a reduction in mean error and increase in Lin’s Concordance Correlation Coefficient (CCC) for both cortical and subcortical grey matter. Full details of volumetric analysis can be viewed in supplementary Figxure S4.

**Figure 3).**
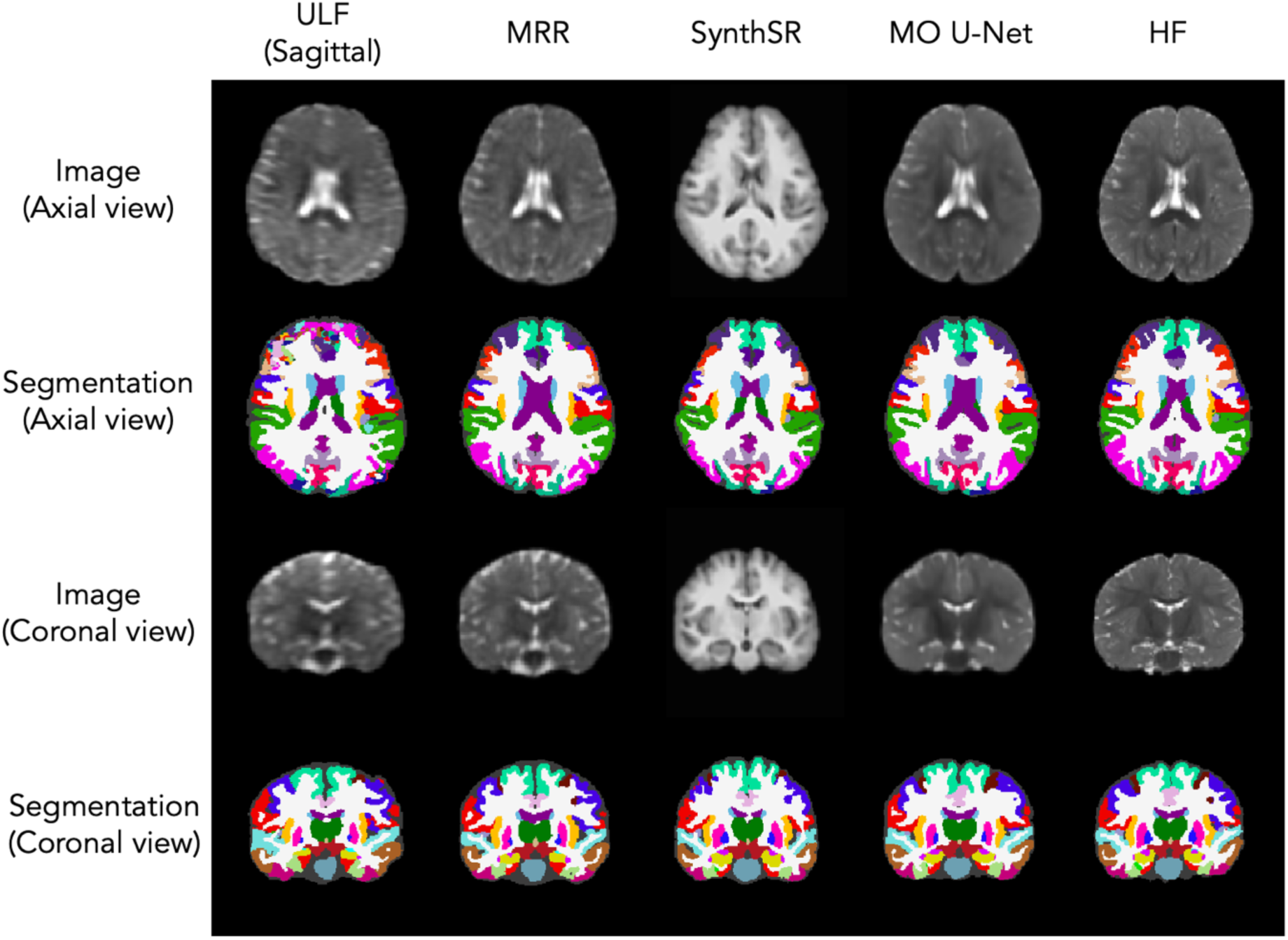
Input images (rows 1 & 3) and corresponding SynthSeg segmentations (rows 2 & 4) from a single test subject. Columns, from left to right: sagittal ULF scan, MRR output, SynthSR output, MO U-Net output, ground-truth HF scan. **Note**: the figure displays SynthSeg outputs including cortical parcellation, however these were merged into single a cortical label (cortical grey matter) for segmentation-based analyses.

**Figure 4).**
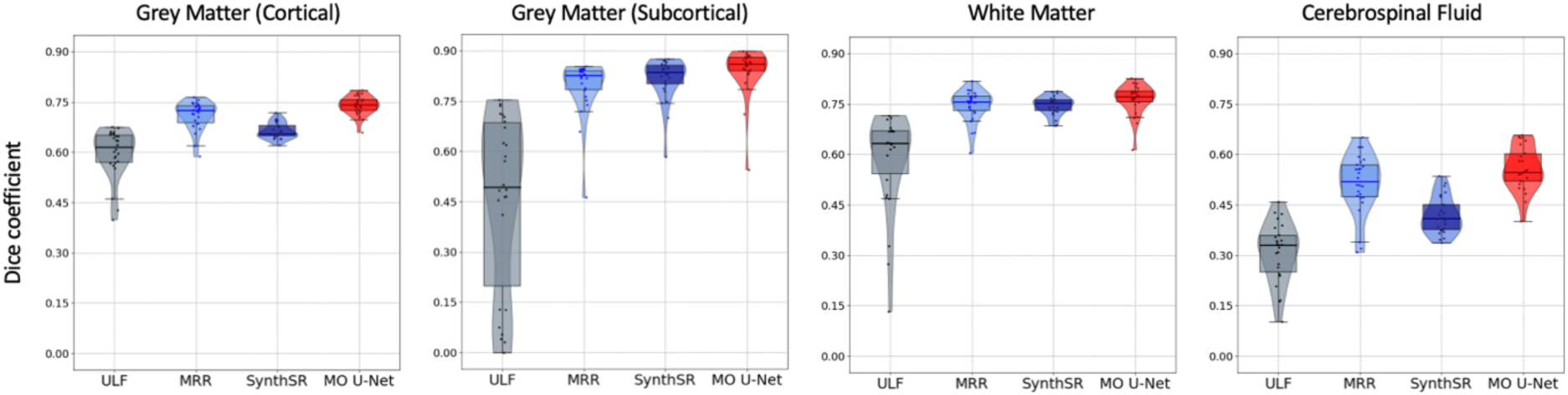
Dice coefficients between HF scan and the original Hyperfine scan (average of axial, coronal, and sagittal), MRR output, SynthSR output, and MO U-Net output, across tissue types. From left to right: cortical grey matter, subcortical grey matter, white matter and cerebrospinal fluid.

**Table 1).** Wilcoxon signed-rank test applied to Dice scores of 26 test subjects, comparing outputs of MO U-Net to MRR, and MO U-Net to SynthSR. For each comparison, the rank biserial correlation (RBC) and associated significance of the underlying Wilcoxon signed-rank test are displayed. Analysis is centred on global tissue types.

| Region | MO U-Net > MRR |  | MO U-Net > SynthSR |  |
| --- | --- | --- | --- | --- |
|  | RBC | Sig. | RBC | Sig. |
| Grey Matter (cortical) | 0.778 | 0.00011* | 0.978 | <0.0001* |
| Grey Matter (subcortical) | 0.852 | <0.0001* | 0.493 | 0.014 |
| White Matter | 0.556 | 0.0060* | 0.652 | 0.0013* |
| Cerebrospinal Fluid | 0.704 | 0.00052* | 0.994 | <0.0001* |

When examining individual subcortical regions, we similarly observed increased Dice overlap of MO U-Net predictions compared to ULF scans across all structures. The same pattern was seen in comparison with MRR outputs, and in five out of eight regions in comparison to SynthSR outputs. For brevity, the Dice scores of subcortical regions showing minimum, median, and maximum MO U-Net performance are depicted in Figure 5, however a full summary table can be found in supplementary Table S5, with significance testing in Table S6 and age stratification in Table S7. When examining subcortical volumes, we observed that the MO U-Net produced outputs with similar linear correlation to high-field volumes as other SR techniques, however we found that the MO U-Net deviated from ground-truth subcortical volumes by the smallest margin. This is evidenced by a mean difference between high-field and predicted volumes (across regions) of −0.281cm^3^, compared to −0.521cm^3^ and 0.506cm^3^ for MRR and SynthSR, respectively, and mean Lin’s CCC of 0.73, compared to 0.66 and 0.57 for MRR and SynthSR, respectively. Subcortical regions with minimum, median, and maximum correlation values are depicted in Figure 6, with a full summary provided in supplementary Table S8.

**Figure 5).**
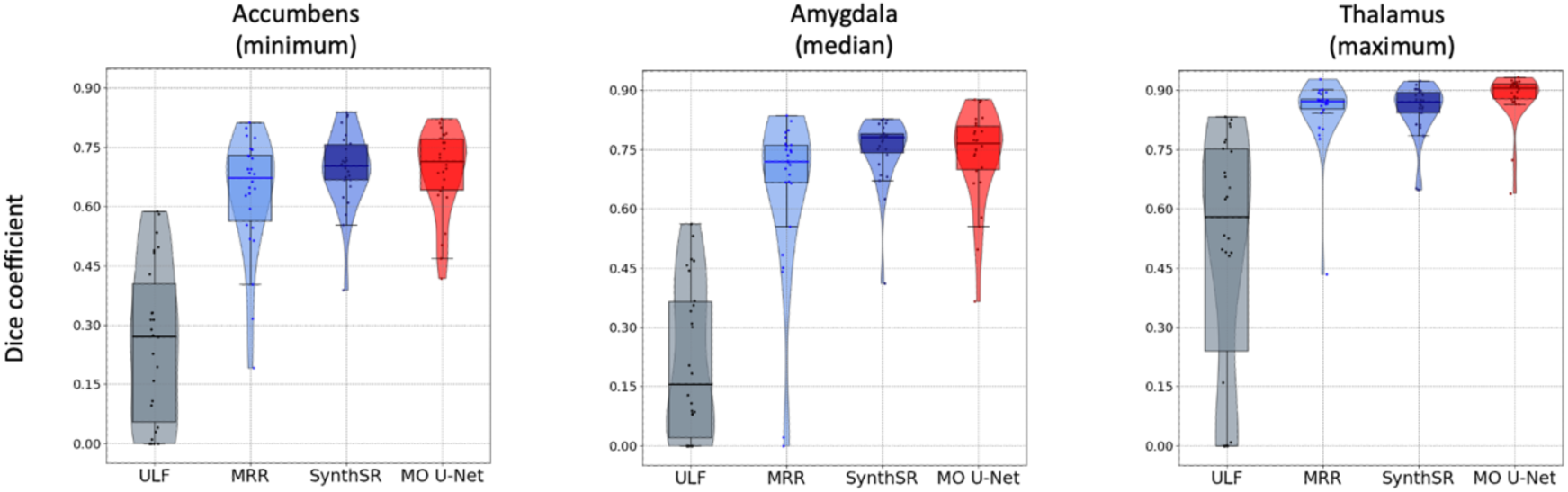
Dice coefficients between HF scan and the original ULF scan (average of axial, coronal, and sagittal), MRR output, SynthSR output, and MO U-Net output, across subcortical regions. From left to right: accumbens (minimum), amygdala (median) and thalamus (maximum).

**Figure 6).**
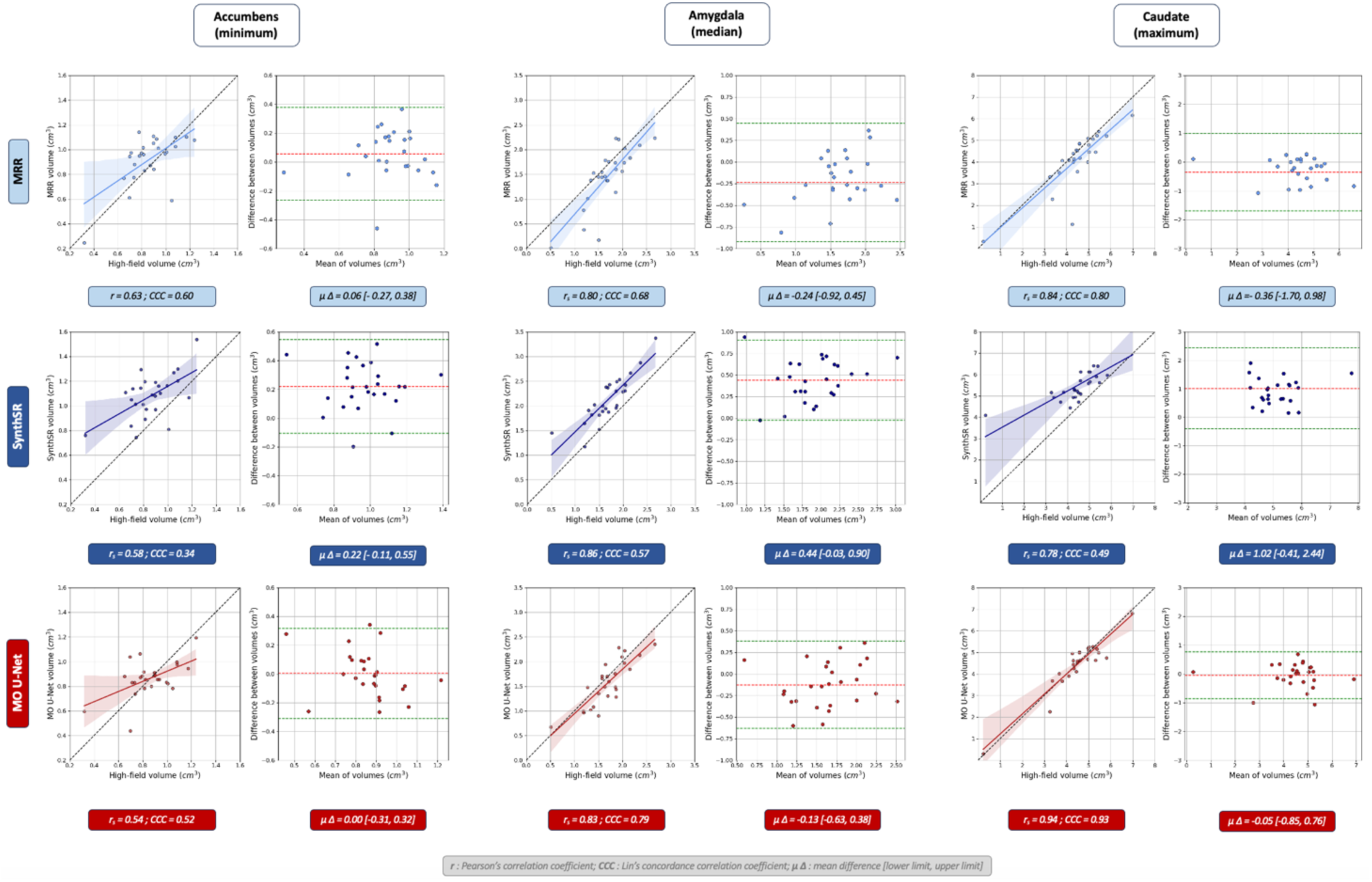
Region-specific volume analysis between MRR outputs and ground-truth HF scans (light blue), SynthSR outputs and ground-truth HF-scans (dark blue), and MO U-Net outputs and ground-truth HF scans (red). For each SR method and region, we display volume correlation (left box) and Bland-Altman plot (right box). Regions, from left to right, include: accumbens (minimum r), amygdala (median r) and caudate (maximum r).

### 3.2 Intensity differentiation and image quality assessment

We next compared the quality of all output T_2_w images. Predictions from the MO U-Net yielded visually superior images compared to both original ULF scans and MRR outputs; MO U-Net outputs appeared less noisy and captured fine details more accurately (Figure 7). The enhanced quality of outputs is further evidenced by greater GM/WM intensity differentiation, with a percentage increase in median difference between GM and WM intensities relative to ULF scans of 39.4% for the MO U-Net compared to 12.7% for MRR (Table 2).

**Figure 7).**
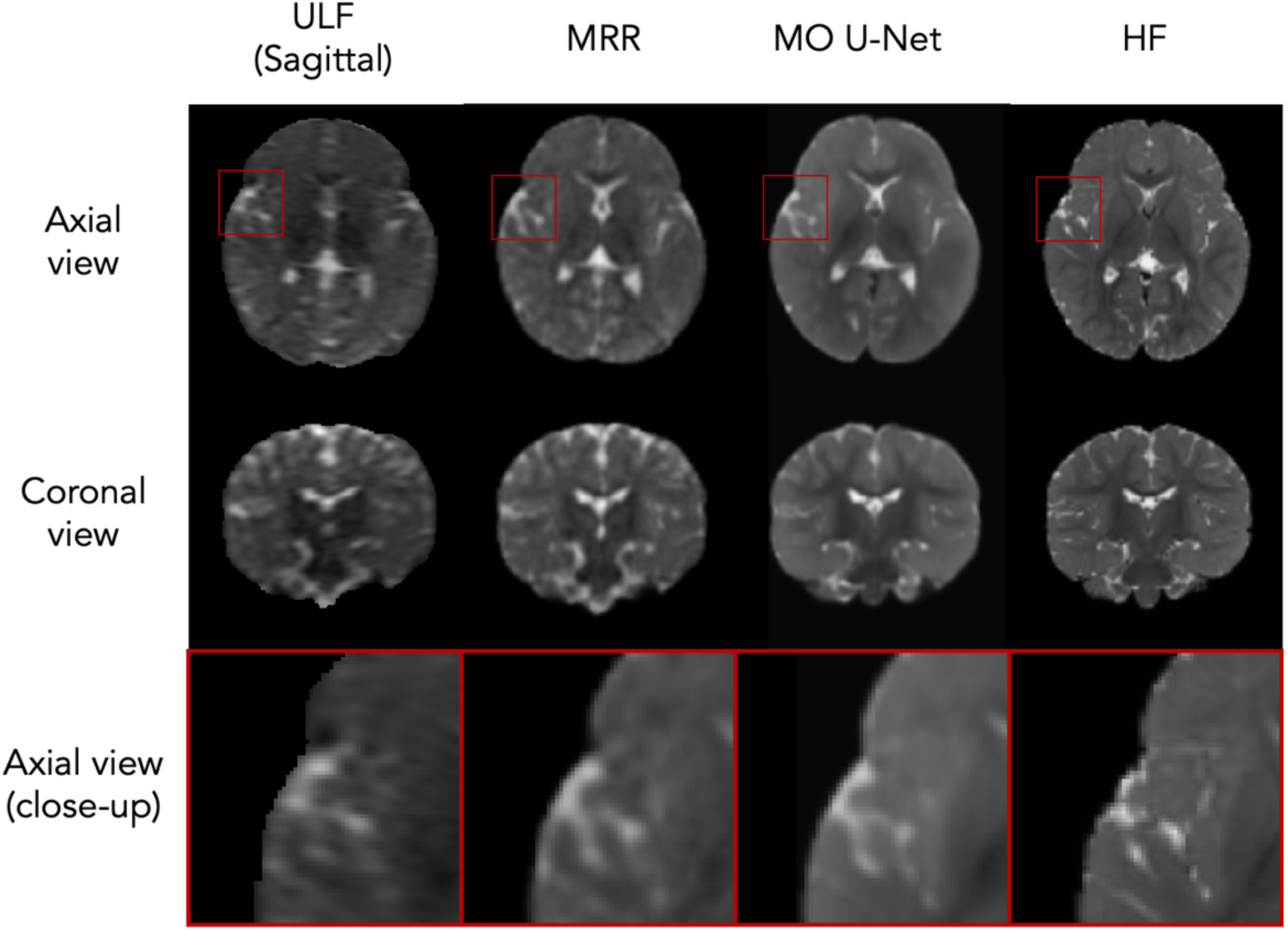
Model outputs from a single test subject. Left to right: raw sagittal ULF scan, MRR output, MO U-Net output and ground-truth HF scan.

**Table 2).** Absolute difference in median grey matter and median white matter intensity in ULF scans, MRR outputs, MO U-Net outputs, and ground-truth high-field scans. The percentage difference in intensity differentiation from ULF ((X-ULF)/ULF) is also shown. The analyses are conducted on N=28 test subjects.

| Intensity measure | ULF (average) | MRR | MO U-Net | HF |
| --- | --- | --- | --- | --- |
| Median difference (GM vs WM) | 0.0213 | 0.0240 | 0.0297 | 0.0285 |
| % difference relative to ULF | 0.00 | $\pm 12.7$ | $\pm 39.4$ | $\pm 33.7$ |

When assessing how well each SR technique’s output matched their corresponding high-field scan (i.e. T_2_w high-field for MRR and MO U-Net vs T_1_w high-field for SynthSR), MO U-Net predictions yielded the lowest NMSE, highest PSNR, and highest SSIM (Table 3). As stated in the Methods section, these analyses were conducted on a subset of N=25 test subjects who had a T_1_w HF scan in addition to their T_2_w HF scan, however analyses on all 28 T_2_w scans for MRR and MO U-Net outputs (excluding SynthSR) yielded nearly identical results (see supplementary Table S9). Additionally, repeating the same analyses after age stratification indicated that our MO U-Net had greater consistency in output quality on 3-month-old subjects and 6-month-old subjects than either MRR or SynthSR (see supplementary Table S10).

**Table 3).** Image quality metrics (NMSE, PSNR, SSIM) for each SR method: MRR, SynthSR and MO U-Net. Values are obtained by comparing SR outputs with ground-truth HF scans. The analyses are conducted on N=25 test subjects.

| SR Method | NMSE (↓) | PSNR (↑) | SSIM (↑) |
| --- | --- | --- | --- |
| MRR | 0.162 | 26.696 | 0.451 |
| SynthSR | 0.742 | 21.377 | 0.872 |
| MO U-Net | **0.063** | **30.814** | **0.906** |

### 3.3 Performance with reduced number of scans

Finally, we assessed how our pre-trained model performed using a reduced number of inputs. We observed that the MO U-Net performed best given all three inputs (axial, coronal and sagittal), with average Dice score rising from 0.723 to 0.730 from one input to all three in global tissue-types (Table 4), and from 0.775 to 791 across individual subcortical regions (supplementary Table S11). Moreover, we found that regardless of the number of distinct input scans used, the MO U-Net yielded the highest Dice scores across all global tissue types, outperforming both SynthSR and MRR even when using only a single axial scan from each subject for inference.

**Table 4).** Tissue-specific Dice scores between HF segmentations and segmentations obtained from ULF scans, MRR outputs, SynthSR outputs and MO U-Net outputs. MO U-Net scores are further stratified according to how many unique inputs the model received (A = axial, A/S = axial + sagittal, A/C/S = axial + sagittal + coronal).

| Region | ULF (avg) | MRR | SynthSR | MO U-Net |  |  |
| --- | --- | --- | --- | --- | --- | --- |
|  |  |  |  | A | A/S | A/C/S |
| GMC | 0.615 | 0.725 | 0.655 | 0.732 | **0.742** | 0.741 |
| GMS | 0.493 | 0.827 | 0.836 | 0.842 | 0.857 | **0.861** |
| WM | 0.633 | 0.757 | 0.753 | 0.765 | 0.768 | **0.772** |
| CSF | 0.331 | 0.520 | 0.409 | **0.554** | 0.535 | 0.547 |
| Average | 0.517 | 0.707 | 0.663 | 0.723 | 0.726 | **0.730** |

## 4 Discussion

Obtaining accurate, high-resolution MRI scans in paediatric populations is essential for studies of neurodevelopment and for diagnosis of neurological conditions. In LMICs where MRI accessibility is significantly reduced, this may be achieved via the use of ULF scanners, supplemented with techniques to improve scan quality. Here we demonstrate improved results compared to previous efforts (Deoni et al., 2022; Iglesias et al., 2023) using our MO U-Net for SR of ULF paediatric images. We present greater Dice overlap of segmentations and increased agreement of corresponding volumes across most brain regions, as well as improved scores on quality metrics of underlying images.

By setting the MO U-Net to have three input channels for each of three separate ULF scans, our model is designed to maximise the amount of anatomical data available from a single subject. As such, our model shows peak performance when all three inputs are provided, however Dice scores from model outputs with varying numbers of unique input scans indicate that a single input is sufficient to produce outputs that outperform other techniques. Tolerance to long scanning times is low in infants (Barkovich et al., 2019), thus successfully completing three separate 3-6 minute-scanning sessions is often infeasible. As such, having a model that can reconstruct an image from just one ULF scan has major practical benefits. An alternative approach would involve modifying our architecture to have one full encoding branch for each input image, where these separate encodings are then combined before they enter the expanding path (Lau et al 2023). This would encourage distinct, image-specific encodings, which may further maximise the information extracted from each scan, however it would impede the applicability of the model to scenarios with a reduced number of scans. A further alternative to our current method would be to utilise multiple contrasts as opposed to multiple orientations (i.e. to train a model using axial T_1_w and axial T_2_w ULF scans as opposed to axial, coronal and sagittal T_2_w ULF scans), as differing contrast may afford a greater benefit. In our study we opted for multiple orientations, as this allowed for a straightforward comparison with MRR and allowed us to make the most use of a dataset where a large portion of subjects completed three T_2_w ULF scans and comparatively few completed a T_1_w ULF scan (see supplementary Table S12). Finally, models could be trained on single-orientation input scans. Both multi-contrast and single-scan models will be a compelling avenue for further work.

Given the demographic of the dataset we used, our model manages to tackle a particularly difficult image translation problem. Owing to differing water and fat content in the developing brain compared to adults, alterations in signal intensities in newborns and infants are seen with both T_1_- and T_2_-weighted anatomical MRI. In particular, the first 6 months of life are characterised by a reversal of the normal adult contrast (T_1_-weighted: lower WM intensity than GM; T_2_-weighted: higher WM intensity than GM) (Dubois et al., 2021), resulting in an “isointense” phase at the 6-month mark where the intensity distributions of GM and WM show strong overlapping (Bui et al., 2020). These neurodevelopmental changes amplify the drastic difference in contrast between HF and ULF MRI, already present in both T_1_w and T_2_w scans. Despite these additional challenges compared to adolescent or adult MRI datasets, we succeed in producing output images with significantly enhanced quality, where GM/WM differentiation closely resembles that of 3T HF scans.

The lack of contrast in paediatric ULF MRI scans additionally complicates segmentation-based analyses. We used SynthSeg (Billot et al., 2023) as it is the only widely available toolkit agnostic to contrast and resolution, allowing direct comparison with non-isotropic T_2_w ULF scans, 1.5mm T_2_w isotropic MRR outputs, as well as 1mm isotropic T_1_w SynthSR outputs. Moreover, SynthSeg has shown excellent performance on ultra-low-field scans of healthy adults (Váša et al., 2024). However, as SynthSeg was trained on adult data, its application to paediatric images runs the risk of producing unexpected outputs. In particular, we observe that Dice scores from 3-month-old subjects are consistently lower than those of 6-month-old subjects, across all regions tested. This may be due to our MO U-Net being trained primarily on 6-month data, however seeing as this pattern is not replicated when viewing image quality metrics that do not rely on a segmentation (NMSE, PSNR, SSIM), it may rather be an effect of inconsistent segmentations by SynthSeg on an age-group whose brain scans differ significantly from those of adults. Consequently, a more accurate approach to testing our model performance would involve the use of a segmentation model trained specifically on paediatric scans. Given such a reliable segmentation scheme, we could further expand upon our performance metrics with additional boundary-based measures (such as the Hausdorff distance or average symmetric surface distance; Reinke et al., 2024). Nevertheless, we demonstrate that the outputs of our MO U-Net result in higher-quality segmentations of all four global tissue types, and most subcortical regions, than either MRR or SynthSR.

We note that volumes derived from segmentations of all super-resolved outputs show deviations from reference standard estimates derived from HF scans. Both MRR and our MO U-Net tend to underestimate the volumes of GM and WM, including individual subcortical regions, and overestimate the volume of CSF (in line with recent work on adults; Váša et al., 2024). Conversely, SynthSR tends to underestimate cortical GM volumes and overestimate subcortical GM volumes. However, volume estimates derived from our MO U-Net show smaller deviations than other models in most evaluated regions. Multiple factors may contribute to deviations of super-resolved volume estimates from HF scans, as well as differences in these deviations across models. Both deep-learning-based models were trained on fundamentally different data (empirical paediatric scans for our MO U-Net vs synthetic adult scans for SynthSR), and across all models, differences in output image contrast are shown (T_2_w for MRR and MO U-Net vs T_1_w for SynthSR), which may in turn impact partial-volume effects at tissue boundaries.

A limitation of our study is that we only assess model performance on unseen subjects of the same age and from the same scanning site. To fully assess the limits of our model, we would run inference on ULF scans from subjects of varying ages and alternative sites, investigating the magnitude of deviation from the training set that results in a sharp decline in model performance. Additionally, including test subjects with pre-specified neurobiological conditions/lesions would allow us to investigate whether using deep learning-based SR runs the risk of failing to capture pathologies. This would provide insight as to whether a model trained on normative images can be applied to subjects with potential pathologies, or if disease-specific models need to be trained to ensure reliable SR outputs.

Furthermore, although the U-Net has proven to be the gold-standard for many deep-learning applications to medical imaging (Kelly et al., 2022), the use of novel architectures such as diffusion models or vision transformers may yield further improvements in quality of model outputs. These methods have the potential to address critical limitations of previous techniques, further enhancing overall realism in super-resolved images. Within the domain of various medical image translation tasks, both diffusion models and vision transformers have been shown to outperform CNN and GAN-based models in terms of SSIM and PSNR (Dalmaz et al., 2021; Kim and Park, 2023). As such, future work will explore the use of such models.

## Conclusion

Ultra-low-field (ULF) imaging presents a potential paradigm-shift in neuroimaging, however its ingress into widespread research and clinical use is impeded by limitations on image resolution and quality. Here we demonstrate how the use of deep learning can aid in deriving higher-resolution T2-weighted scans of healthy paediatric subjects from ULF scanners. We do so by training our MO U-Net on paired ULF-HF scans using a combined voxel-wise and perceptual loss, and demonstrate superior performance in comparison to alternative deep learning and non-deep learning based methods for super-resolving paediatric scans.

## Code availability statement

All model training code, along with model weights, is available at: https://github.com/levente-1/MO-U-Net.

## Supplementary Information

**Figure S3).**
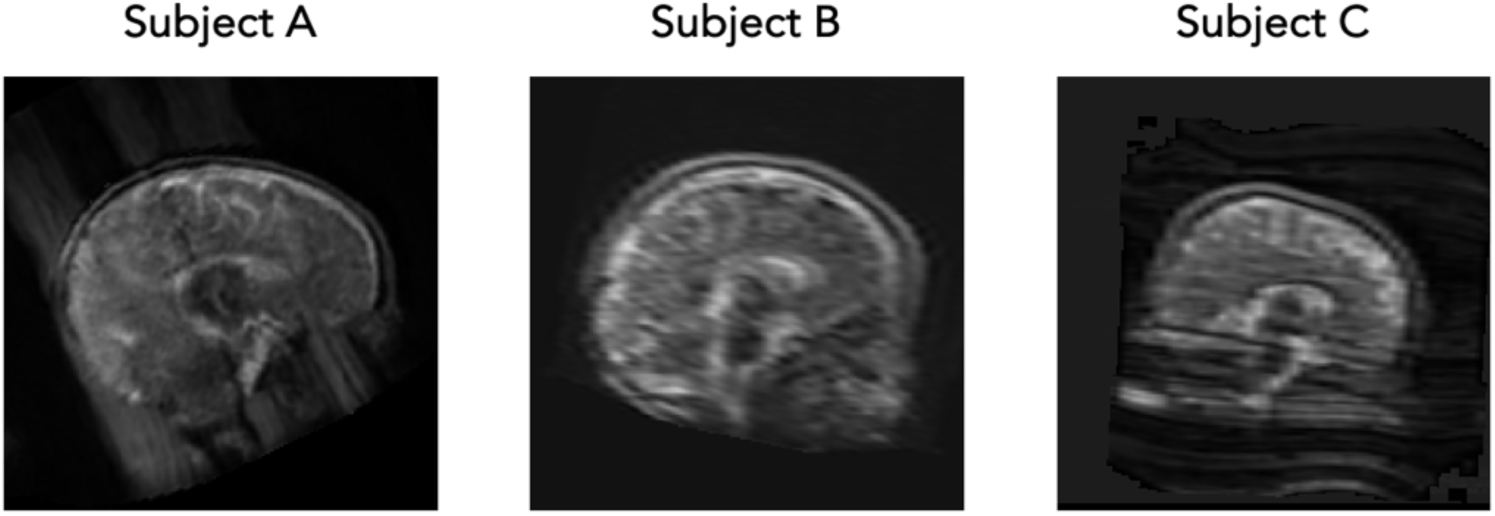
Sample of subjects who were excluded from the training set. All subjects A, B and C exhibit significant imaging artifacts that make them unsuitable for model training or inference.

**Table S2).** Demographic distribution across training, validation and test sets across all four folds.

|  | Age<br>(months) | Training set |  | Validation set |  | Test set |  |
| --- | --- | --- | --- | --- | --- | --- | --- |
|  |  | Male | Female | Male | Female | Male | Female |
| Fold 1 | 3 | 6 (14.3%) | 6 (14.3%) | 1 (14.3%) | 1 (14.3%) | 0 (0.0%) | 1 (14.3%) |
|  | 6 | 17 (40.5%) | 13 (30.9%) | 3 (42.8%) | 2 (28.6%) | 3 (42.8%) | 3 (42.8%) |
| Fold 2 | 3 | 5 (11.9%) | 5 (11.9%) | 1 (14.3%) | 2 (28.6%) | 1 (14.3%) | 1 (14.3%) |
|  | 6 | 19 (45.3%) | 13 (30.9%) | 2 (28.6%) | 2 (28.6%) | 2 (28.6%) | 3 (42.8%) |
| Fold 3 | 3 | 5 (11.9%) | 6 (14.3%) | 1 (14.3%) | 1 (14.3%) | 1 (14.3%) | 1 (14.3%) |
|  | 6 | 17 (40.5%) | 14 (33.3%) | 3 (42.8%) | 2 (28.6%) | 3 (42.8%) | 2 (28.6%) |
| Fold 4 | 3 | 5 (11.9%) | 6 (14.3%) | 1 (14.3%) | 1 (14.3%) | 1 (14.3%) | 1 (14.3%) |
|  | 6 | 16 (38.1%) | 15 (35.7%) | 4 (57.1%) | 1 (14.3%) | 3 (42.8%) | 2 (28.6%) |

**Table S2).**
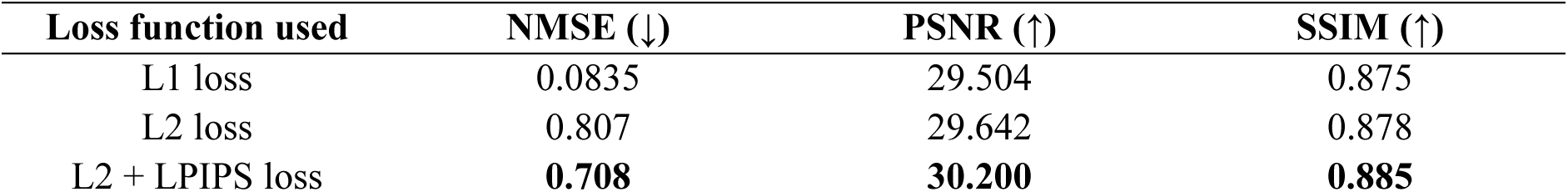
Image quality assessment of predictions from MO U-Nets trained on varying loss functions (L1, L2, L2 + LPIPS). All models were trained for 1000 epochs on a subset of the data used in the final experiment.

| Loss function used | NMSE (↓) | PSNR (↑) | SSIM (↑) |
| --- | --- | --- | --- |
| L1 loss | 0.0835 | 29.504 | 0.875 |
| L2 loss | 0.807 | 29.642 | 0.878 |
| L2 + LPIPS loss | **0.708** | **30.200** | **0.885** |

**Figure S2).**
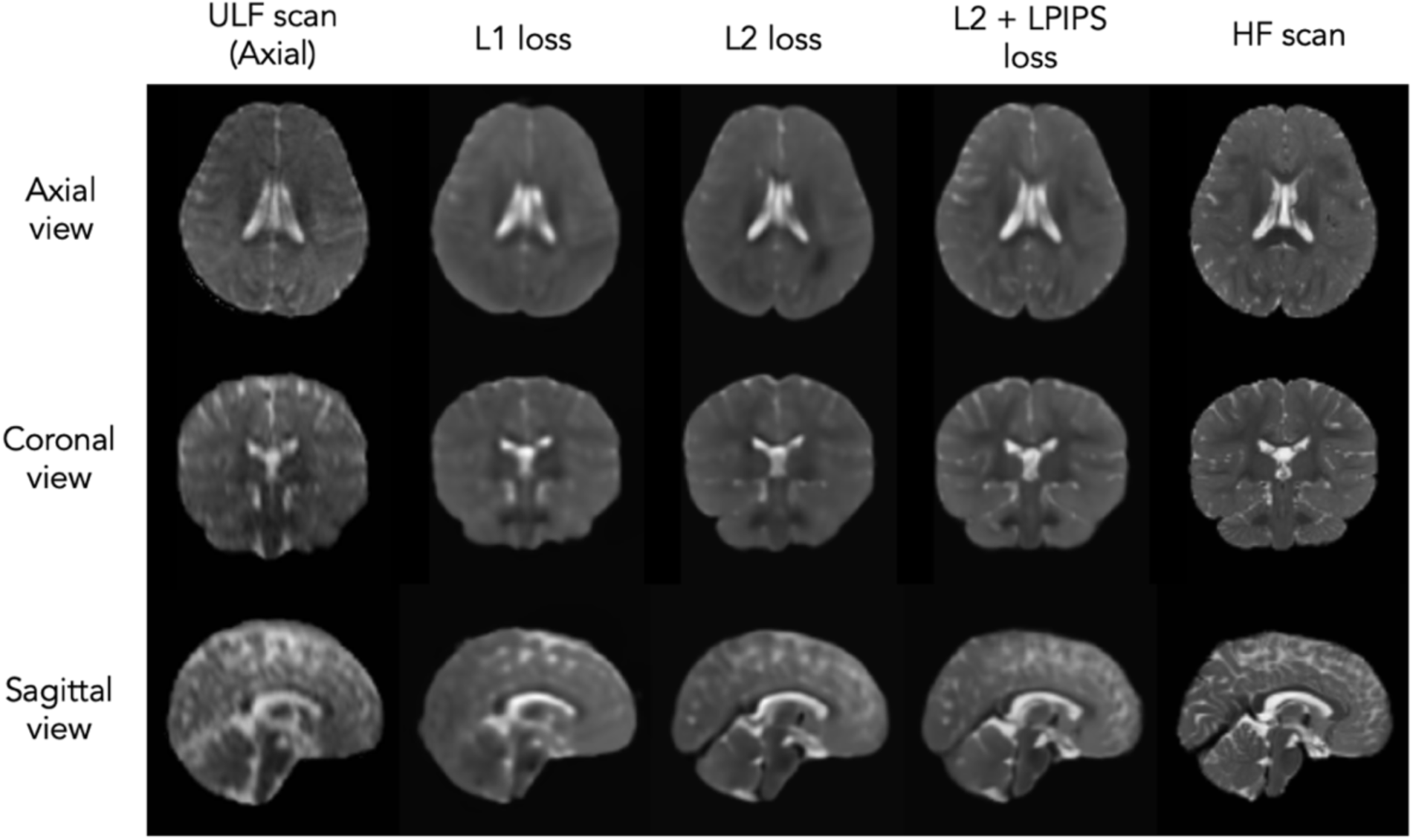
Model outputs from MO U-Nets trained on varying loss functions. Left to right: axial ULF scan, MO U-Net output with L1 loss, L2 loss, L2 + LPIPS loss, ground-truth HF scan.

**Figure S3).**
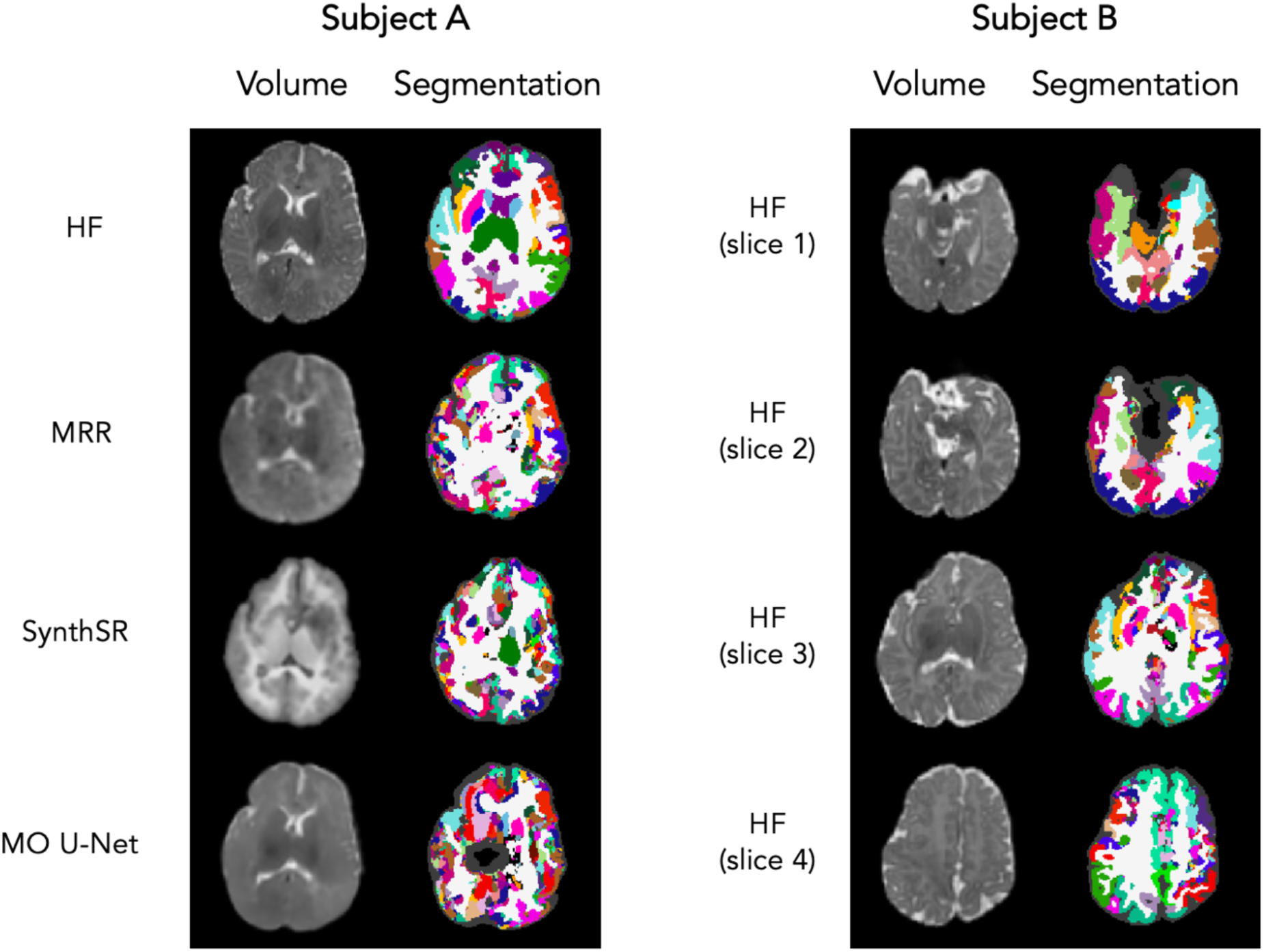
Test subjects excluded from segmentation-based analyses, owing to poor SynthSeg outputs. A) Subject with acceptable HF segmentation, but failed segmentations for all three SR outputs (MRR, SynthSR, and MO U-Net). B) Subject with failed HF segmentation, preventing comparison to ground-truth. Both subjects excluded had scans taken at 3-months of age.

**Table S3).**
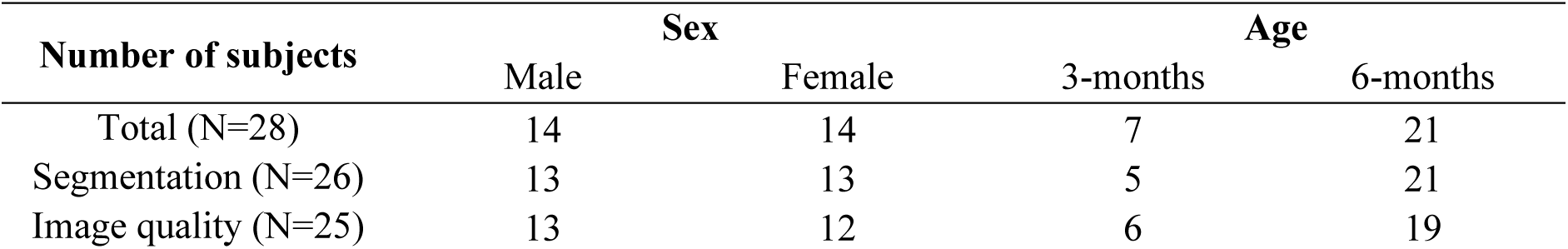
Distribution of sex and age across test set. Row 1: total test subjects; Row 2: test subjects for segmentation-based analyses. where two subjects with failed segmentations were excluded; Row 3: test subjects for image quality metrics (NMSE, PSNR, SSIM), where three subjects with no T_1_w HF scan were excluded.

**Table S4).** Median Dice overlap with segmentations from HF scans, across global tissue types. Scores are stratified according to age group: 3-months (N=5) and 6-months (N=21). Both values are shown for ULF scans, MRR outputs, SynthSR outputs and MO U-Net outputs.

| Region | ULF (avg) |  | MRR |  | SynthSR |  | MO U-Net |  |
| --- | --- | --- | --- | --- | --- | --- | --- | --- |
|  | 3M | 6M | 3M | 6M | 3M | 6M | 3M | 6M |
| GMC | 0.553 | 0.635 | 0.726 | 0.719 | 0.657 | 0.654 | 0.726 | 0.742 |
| GMS | 0.054 | 0.585 | 0.752 | 0.828 | 0.748 | 0.847 | 0.843 | 0.864 |
| WM | 0.469 | 0.633 | 0.734 | 0.760 | 0.718 | 0.753 | 0.708 | 0.776 |
| CSF | 0.208 | 0.332 | 0.530 | 0.509 | 0.427 | 0.408 | 0.520 | 0.548 |
| **Average** | 0.321 | **0.546** | 0.686 | **0.704** | 0.638 | **0.666** | 0.699 | **0.732** |

**Figure S4).**
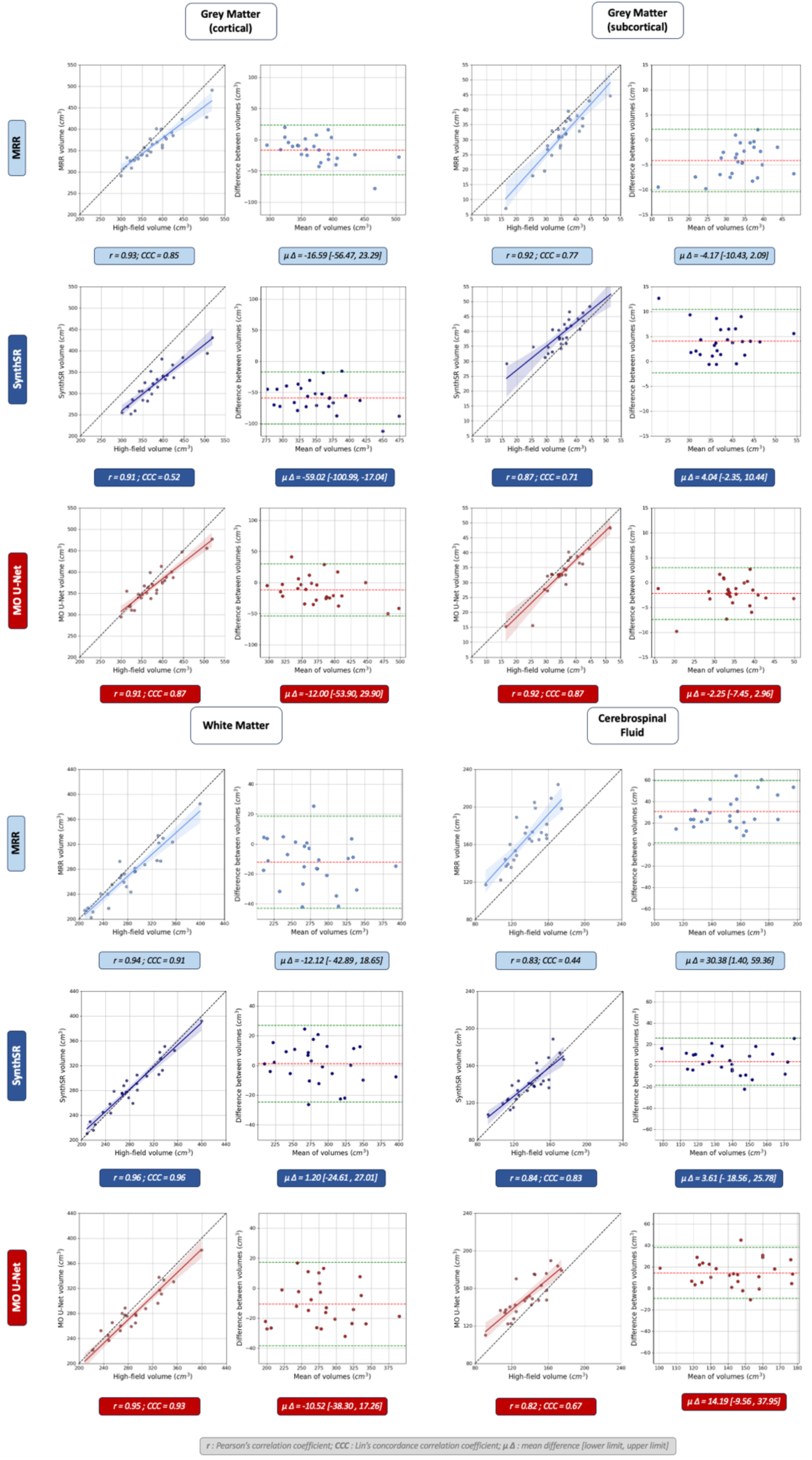
Tissue-type-specific volume analysis between MRR outputs and ground-truth HF scans (light blue), SynthSR outputs and ground-truth HF-scans (dark blue), and MO U-Net outputs and ground-truth HF scans (red). For each SR method and tissue type, we display volume correlation (left) and Bland-Altman plot (right). Tissue types, from left to right, top to bottom: cortical grey matter, subcortical grey matter, white matter, cerebrospinal fluid.

**Table S5).** Median Dice overlap with segmentations from HF scans, across subcortical regions. Columns 2-5: average of ULF scans (axial, coronal and sagittal), MRR outputs, SynthSR outputs, MO U-Net outputs.

| Region | ULF (avg) | MRR | SynthSR | MO U-Net |
| --- | --- | --- | --- | --- |
| Accumbens | 0.272 | 0.673 | 0.703 | 0.715 |
| Amygdala | 0.156 | 0.720 | 0.782 | 0.766 |
| Caudate | 0.412 | 0.747 | 0.735 | 0.803 |
| Hippocampus | 0.256 | 0.698 | 0.708 | 0.764 |
| Pallidum | 0.289 | 0.625 | 0.799 | 0.741 |
| Putamen | 0.456 | 0.781 | 0.819 | 0.838 |
| Thalamus | 0.579 | 0.872 | 0.870 | 0.906 |
| Ventral DC | 0.374 | 0.786 | 0.808 | 0.798 |
| **Average** | **0.349** | **0.738** | **0.778** | **0.791** |

**Table S6).** Wilcoxon signed-rank test applied to Dice scores of 26 test subjects, comparing outputs of MO U-Net to MRR, and MO U-Net to SynthSR. For each comparison, the rank biserial correlation (RBC) and associated significance of the underlying Wilcoxon signed-rank test are displayed. Analysis is centred on subcortical regions. FWR = 0.003125.

| Region | MO U-Net > MRR |  | MO U-Net > SynthSR |  |
| --- | --- | --- | --- | --- |
|  | RBC | Sig. | RBC | Sig. |
| Accumbens | 0.385 | 0.045 | 0.569 | 0.57 |
| Amygdala | 0.556 | 0.0060 | 0.037 | 0.44 |
| Caudate | 0.749 | 0.00021* | 0.442 | 0.025 |
| Hippocampus | 0.943 | <0.0001* | 0.601 | 0.0031 |
| Pallidum | 0.778 | 0.00011* | -0.413 | 0.97 |
| Putamen | 0.875 | <0.0001* | 0.162 | 0.24 |
| Thalamus | 0.726 | 0.00033* | 0.675 | 0.00089* |
| Ventral DC | 0.527 | 0.0088 | 0.265 | 0.12 |

**Table S7).** Median Dice overlap with segmentations from HF scans, across subcortical regions. Scores are stratified according to age group: 3-months (N=5) and 6-months (N=21). Both values are shown for ULF scans, MRR outputs, SynthSR outputs and MO U-Net outputs.

| Region | ULF (avg) |  | MRR |  | SynthSR |  | MO U-Net |  |
| --- | --- | --- | --- | --- | --- | --- | --- | --- |
|  | 3M | 6M | 3M | 6M | 3M | 6M | 3M | 6M |
| Accumbens | 0.010 | 0.313 | 0.518 | 0.694 | 0.667 | 0.716 | 0.648 | 0.723 |
| Amygdala | 0.0 | 0.301 | 0.441 | 0.721 | 0.671 | 0.783 | 0.664 | 0.774 |
| Caudate | 0.006 | 0.445 | 0.679 | 0.771 | 0.646 | 0.764 | 0.781 | 0.804 |
| Hippocampus | 0.0 | 0.320 | 0.385 | 0.703 | 0.512 | 0.729 | 0.559 | 0.778 |
| Pallidum | 0.0 | 0.308 | 0.448 | 0.630 | 0.534 | 0.804 | 0.675 | 0.754 |
| Putamen | 0.010 | 0.515 | 0.690 | 0.781 | 0.735 | 0.833 | 0.804 | 0.841 |
| Thalamus | 0.0 | 0.654 | 0.806 | 0.875 | 0.806 | 0.876 | 0.904 | 0.910 |
| Ventral DC | 0.0 | 0.432 | 0.701 | 0.793 | 0.711 | 0.815 | 0.781 | 0.800 |
| **Average** | **0.003** | **0.411** | **0.584** | **0.746** | **0.660** | **0.790** | **0.727** | **0.798** |

**Table S8).**
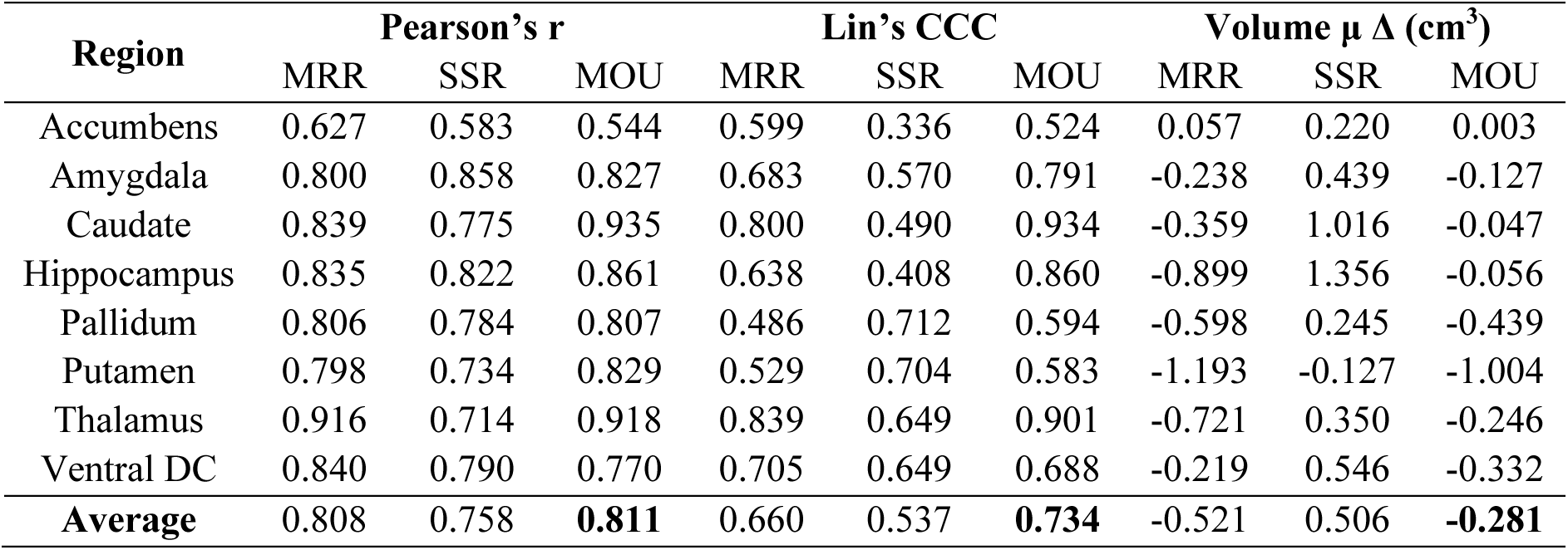
Pearson’s r, Lin’s CCC, and mean estimated volume difference as calculated for each SR technique. MRR = multi-resolution registration, SSR = SynthSR, MOU = multi-orientation U-Net.

| Region | Pearson's $r$ | | | Lin's CCC | | | Volume $\mu \Delta$ (cm<sup>3</sup>) | | |
| --- | --- | --- | --- | --- | --- | --- | --- | --- | --- |
|  | MRR | SSR | MOU | MRR | SSR | MOU | MRR | SSR | MOU |
| Accumbens | 0.627 | 0.583 | 0.544 | 0.599 | 0.336 | 0.524 | 0.057 | 0.220 | 0.003 |
| Amygdala | 0.800 | 0.858 | 0.827 | 0.683 | 0.570 | 0.791 | -0.238 | 0.439 | -0.127 |
| Caudate | 0.839 | 0.775 | 0.935 | 0.800 | 0.490 | 0.934 | -0.359 | 1.016 | -0.047 |
| Hippocampus | 0.835 | 0.822 | 0.861 | 0.638 | 0.408 | 0.860 | -0.899 | 1.356 | -0.056 |
| Pallidum | 0.806 | 0.784 | 0.807 | 0.486 | 0.712 | 0.594 | -0.598 | 0.245 | -0.439 |
| Putamen | 0.798 | 0.734 | 0.829 | 0.529 | 0.704 | 0.583 | -1.193 | -0.127 | -1.004 |
| Thalamus | 0.916 | 0.714 | 0.918 | 0.839 | 0.649 | 0.901 | -0.721 | 0.350 | -0.246 |
| Ventral DC | 0.840 | 0.790 | 0.770 | 0.705 | 0.649 | 0.688 | -0.219 | 0.546 | -0.332 |
| **Average** | 0.808 | 0.758 | **0.811** | 0.660 | 0.537 | **0.734** | -0.521 | 0.506 | **-0.281** |

**Table S9).** Image quality metrics (NMSE, PSNR, SSIM) for SR methods generating T_2_w scans: MRR and MO U-Net. Values are obtained by comparing SR outputs with ground-truth HF scans. The analyses are conducted on all N=28 test subjects.

| SR Method | NMSE (↓) | PSNR (↑) | SSIM (↑) |
| --- | --- | --- | --- |
| MRR | 0.166 | 26.164 | 0.447 |
| MO U-Net | **0.068** | **30.527** | **0.901** |

**Table S10).** Image quality metrics (NMSE, PSNR, SSIM) for each SR method: MRR, SynthSR and MO U-Net. Scores are stratified according to age group: 3-months (N=6) and 6-months (N=19)

| SR Method | NMSE (↓) | | PSNR (↑) | | SSIM (↑) | |
| --- | --- | --- | --- | --- | --- | --- |
|  | 3-month | 6-month | 3-month | 6-month | 3-month | 6-month |
| MRR | **0.126** | 0.175 | **27.770** | 26.357 | **0.494** | 0.438 |
| SynthSR | **0.277** | 0.954 | **24.032** | 20.539 | **0.878** | 0.870 |
| MO U-Net | **0.063** | **0.063** | **30.826** | 30.810 | 0.875 | **0.915** |

**Table S11).** Dice scores between HF segmentations and segmentations obtained from MO U-Net outputs, across subcortical regions. MO U-Net scores are further stratified according to how many unique inputs the model received (A = axial, A/S = axial + sagittal, A/C/S = axial + sagittal + coronal). For comparison with ULF scans, MRR outputs and SynthSR outputs, see Table S5.

| Region | MO U-Net |  |  |
| --- | --- | --- | --- |
|  | Axial | Axial/Sagittal | Axial/Coronal/Sagittal |
| Accumbens | 0.707 | 0.717 | 0.715 |
| Amygdala | 0.724 | 0.765 | 0.766 |
| Caudate | 0.813 | 0.808 | 0.803 |
| Hippocampus | 0.738 | 0.744 | 0.764 |
| Pallidum | 0.713 | 0.728 | 0.741 |
| Putamen | 0.818 | 0.829 | 0.838 |
| Thalamus | 0.898 | 0.896 | 0.906 |
| Ventral DC | 0.785 | 0.801 | 0.798 |
| **Average** | 0.775 | 0.786 | **0.791** |

**Table S12).**
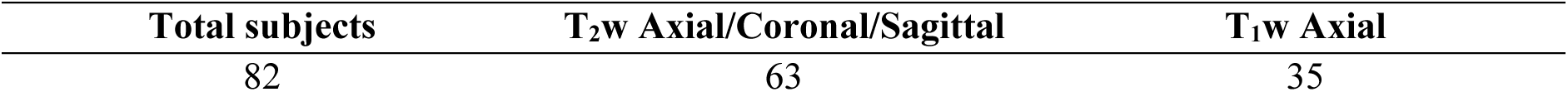
Portion of total subjects having completed three separate T_2_w ULF scans (axial, coronal and sagittal), compared to those who additionally completed a T_1_w axial scan.

## Khula SA Study Team

Michal R. Zieff

Donna Herr

Chloë A. Jacobs

Sadeeka Williams

Zamazimba Madi

Nwabisa Mlandu

Tembeka Mhlakwaphalwa

Lauren Davel

Reese Samuels

Zayaan Goolam

Thandeka Mazubane

Bokang Methola

Khanyisa Nkubungu

Candice Knipe

